# Comparative analysis of adhesin-like proteins from *Methanobacteriales* species

**DOI:** 10.64898/2026.08.03.742440

**Authors:** Abel Tan, Anjali Bansal Gupta, Shuan Er, Henning Seedorf

## Abstract

Methanogenic archaea are key syntrophic partners in animal and human guts. However, the molecular basis of physical and functional interactions of methanogens with other cells remains poorly understood. Adhesin-like proteins (ALPs) are abundant, unusually large, and repeat-rich surface proteins and are potential prime candidates for mediating cell–cell interactions, however, due to poor characterization, ALPs often remain neglected in annotations of genomes and large-scale datasets. Here, we combined structure-guided analysis with comparative genomics to map ALP domain architectures across *Methanobacteriales* species, with emphasis on intestinal and rumen lineages. We curated predicted ALP structures and manually delineated domain boundaries to build a reference set encompassing ABD, RBH, membrane-anchoring domains (MAD), and miscellaneous accessory domains. Leveraging on a novel approach for ALP domain annotations, we generate ALP annotations and domain architectures for 17 *Methanobacteriales* species without reliance on primary-sequence homology. Applying this approach to strains of *Methanobrevibacter smithii* and *Methanobrevibacter intestini*, we assembled species-level “adhesiomes” and assessed conservation patterns of ALPs. Network analysis indicated that ALPs are more commonly shared among strains within a species than between species, suggesting species-specific adhesin repertoires. While total ALP counts differed significantly between *M. smithii* and *M. intestini* (Wilcoxon rank-sum p = 0.00295), median ALP lengths did not (p = 0.606). Our results provide a comprehensive overview of the ALP domain architectures in *Methanobacteriales* species and reveals species-specific adhesiomes that likely contribute to niche adaptation and syntrophic partnerships in gut ecosystems.

## Introduction

Methanogenic archaea from the intestinal tract of animals strongly contribute to fermentative activities in various gut environments (1-3). The methanogens rely hereby mostly on the fermentation products of other microorganisms, such as bacteria, fungi and protozoa, which ultimately breakdown complex carbohydrates into CO_2_, hydrogen, acetate, etc (4). It has been shown previously that methanogens can be closely associated with other microorganisms to form syntrophic relationships (5, 6). However, the molecular basis for these interactions remains poorly understood. Adhesin-like proteins (ALPs) may offer in this regard some potential cues. ALPs from methanogenic archaea were first described as “asparagine-threonine-rich proteins” in the genome of *Methanosphaera stadtmanae* (7). With the subsequent discovery of these proteins in the complete genomes of *Methanobrevibacter smithii* and *Mbb. ruminantium* it became apparent that ALPs are widespread in methanogens, particularly those from the rumen and the human gut (8, 9). As the ‘asparagine-threonine-rich proteins’ shared some similarity with bacterial adhesins they were renamed by Samuel et al. to ‘adhesin-like proteins’ (8), a naming that has been used consistently in literature. While ALPs often share partial sequence identity within a genome, it can be more challenging to perform sequence-based comparisons across species or also to reliable identify them. Previously published reports on methanogen genomes have reported varying numbers of ALPs per species (7-10). Comparative analyses of ALPs across methanogen species remain scarce. A previous study of human gut methanogens identified fewer ALPs (11), whereas a recent study provided a comprehensive overview of ALPs in intestinal *Methanobacteriota* (12). Other large collections of human and rumen methanogen genomes assembled from metagenomic sequencing data did not include ALPs in their analyses (13).. Considering that ALPs are significant features in methanogen genomes, e.g. making up ∼10% of the genomes of *Msp. stadtmanae* and *Mbb. smithii* (14), this represents a significant knowledge gap.

Some of the striking characteristics of ALPs are their length, which can in some cases exceed 8000 amino acids, and the highly repetitive sequence motifs. Insight into the secondary and tertiary structure of ALPs have been scarce until recently, due the absence of any structural analysis of these proteins or their domains. AI-assisted protein prediction programs, such as AlphaFold have provided a glimpse into what ALP domains and potentially entire ALPs may look like (14). A recent analysis of the “adhesiome” (collection of all ALPs) from *Msp. stadtmanae* and *Mbb. smithii* revealed that ALPs have domains that appear to be conserved across ALPs within a methanogen, but also across species. However, the sequence similarity of the domains can be low, even within an ALP, and particularly across species (14).

One of the most frequently observed domains in this regard is ABD, named Archaeal BIg Domain in reference to the ‘Bacterial Immunoglobulin-like domain’. The ABD domains are common in ALPs and can reach copy numbers of more than 40 in large ALPs. While these domains can share relatively low sequence similarity (depending also on methanogen species), they do contain conserved residues that may constitute binding motifs and may also contribute to the overall repetitive structure of these proteins. Right-hand-beta helical domains (RBH) are also commonly found in ALPs. They can vary substantially in length and may also be present in different copy numbers, though not as high as ABDs. Both ABD and RBH are most commonly found in ALPs and the presence of at least either one has been proposed as important criterion to determine if a protein may represent an ALP (14). The identification and structural characterization of ALPs and their domains with bioinformatic tools remains difficult due to the high sequence variation between ALPs and the domains within. This is partially also due to the automated annotation of ABDs and ABD repeats by NCBI’s CDD, which annotates such as ClfA, Big_3_5, Duff11, Mg4 and others, while RBH domains may also be annotated as AIDA, FhaB, NosDm, and others. However, most of the respective ABD and RBH represent similar folds as has become apparent from the analysis of the predicted tertiary structures. This study aims therefore on the analysis of the domain architecture of ALPs from a larger repertoire of *Methanobacteriales* genomes - obtained from different regions of the gastrointestinal tract, including the rumen and intestines, while others are environmental. We then apply this approach to obtain a more comprehensive overview of the adhesiome of *Methanobacteriales* species and determine also species-specific differences in the adhesin microbiome using representative strains of *Mbb. smithii* and *Mbb. intestini*. In summary, this analysis provides an approach to automated ALP domain architecture analysis and also insights into the ALP repertoire of *Methanobacteriales*.

## Materials und Methods

### Genomes and ALP identification

Methanogen genomes of 17 methanogen species as well as 12 additional genomes of *Mbb. smithii* and *Mbb. intestini* strains were downloaded from public resources, such as NCBI. Mostly genome sequences with 10 or less contig were selected to ensure that assemblies contain primarily complete ALPs.

Taxonomic and sequence metadata were consolidated with amino acid sequences, sequence lengths, protein annotations, UniProt identifiers where available, AlphaFold model filenames, and internal protein IDs. Database provenance was assigned at the species level based on the source of each protein record. *Methanobacterium formicicum* was represented by GenBank records, whereas *Mbt. lacus, Mbt. paludis*, and *Methanobacterium* sp. MB1 were represented by RefSeq records. *Msp. stadtmanae, Methanosphaera* sp. BMS, and *Methanosphaera* sp. ISO3-F5 were represented by RefSeq records. *Methanothermobacter* and *Methanothermus* entries were also derived from RefSeq. *Methanobrevibacter* species were represented by a combination of GenBank and RefSeq records (see Table S1 for details). ALPs were identified by querying individual genomes with a pre-defined dataset of 516 ALPs as described in reference 14. Sequences not containing either ABD or RBH were examined manually and discarded as non-ALPs.

### Sequence Retrieval and Structural Modeling

Amino acid sequences for the identified ALPs were retrieved from their corresponding databases (GenBank or RefSeq) based on their accession identifiers. Structural predictions were performed using the AlphaFold Server (https://alphafoldserver.com/) (15). Due to the computational constraints of the modelling architecture, a sequence length threshold was applied: ALPs exceeding 5,000 amino acids were excluded from the structural prediction pipeline as they cannot be modelled by the server.

### Characterization of ALP domains

To characterize the domain architecture of newly identified Adhesin-Like Proteins (ALPs) at scale, we developed an automated pipeline that integrates high-fidelity structural prediction with a domain-recognition framework. This approach was necessitated by the extreme length and modularity of ALPs, which often exceed the practical limits of manual annotation and traditional sequence-based search tools.

The identification and classification of domains within these structures were performed using PRISM (Protein Recognition and Integrated Segmentation Model). PRISM operates by interpreting the unique spatial geometry and topological signatures inherent in the protein structure to perform a “one-shot” segmentation and classification (manuscript in preparation). The model was trained to recognize three primary architectural categories: Archaeal Big Domains (ABD), Right-Handed Beta Helical (RBH) domains, and a collective “Others” category comprising various accessory modules.

The model was trained on a curated dataset of initially 865 manually annotated ALPs derived from diverse species. For performance evaluation, the dataset was divided into an 80% training set and a 20% test set. The efficacy of the framework was assessed based on precision, sensitivity (recall), and overall domain-level accuracy. In instances where the automated predictions deviated from the initial manual annotations, the structures underwent expert review; this iterative process allowed for the identification and correction of manual annotation errors.

### Phylogenetic and Structural Analysis of ALPs

Full-length protein sequences of the curated ALPs were aligned using MAFFT under default parameters (16). Phylogenetic relationships were inferred from the resulting alignment using FastTree (17), which utilizes an approximately maximum-likelihood approach; branch lengths represent expected substitutions per site. Taxonomic metadata, including genus and species designations, were retrieved from NCBI protein record headers. Ecological context regarding host habitats (e.g., human gut, rumen, or environmental) was manually compiled through a targeted literature search for each respective species. The final phylogenetic tree and its associated metadata rings were visualized and annotated using the Interactive Tree Of Life (iTOL) (18).

ALP sequences were classified into distinct structural groups based on the spatial arrangement and composition of their modules as described before (14): Group I (Multi-domain): Sequences containing both ABD and RBH domains without additional non-ALP modules. This group is further divided into Subgroup IA, where all RBH domains are positioned N-terminally to all ABD domains, and Subgroup IB, where ABD and RBH domains are interspersed. Group II (Single-module type): Sequences containing only one of the primary domain types, categorized as Subgroup IIA (ABD-only) or Subgroup IIB (RBH-only). Other: Sequences containing additional functional domains alongside ABDs or RBHs.

### ALP clustering

Sequence metadata were first curated and filtered to retain ALPs from the methanogen strains *Mbb. smithii* and *Mbb. intestini* TLL-48-HuF1. To identify homologous, near-identical ALPs while avoiding the computational burden of all-versus-all sequence comparison, ALPs were first clustered by sequence length and then subclustered by global sequence identity.

Sequences were initially partitioned into two length categories: short proteins, defined as <1,000 amino acids, and long proteins, defined as ≥1,000 amino acids. Within each category, a custom range-constrained hierarchical clustering procedure was applied. Sequences were sorted in ascending order of length and greedily assigned to length-based clusters such that the difference between the shortest and longest sequence in each cluster did not exceed 10 amino acids for short proteins or 50 amino acids for long proteins. Global sequence-identity subclustering was then performed independently within each length-based cluster. Pairwise sequence identities were calculated using the Needleman–Wunsch global alignment algorithm implemented in Biopython’s PairwiseAligner, with a scoring scheme of 1.0 for matches, 0.0 for mismatches, −1.0 for gap opening, and −0.5 for gap extension (19). To improve computational efficiency, pairwise comparisons were restricted to sequence pairs differing in length by no more than 5 amino acids. Subclusters were defined as connected components in a graph in which edges connected sequence pairs with ≥98% global sequence identity.

For network analysis two complementary network representations were generated. In the first approach, a hyperedge-style representation was used in which each ALP was represented by a single square positioned at the centroid of the species nodes in which the ALP occurred. In the second approach, a species–species projection network was constructed based on shared ALP identifiers. For each ALP, all pairwise combinations of host species were enumerated and the number of distinct shared ALPs between species pairs was calculated. These counts were used as undirected edge weights between species nodes. To reduce visual clutter, edges with fewer than two shared ALPs were excluded from the final network by default. Node coordinates were generated using a force-directed Fruchterman–Reingold layout applied to the species graph, in which vertices represented species and weighted edges connected species sharing at least one ALP. Layout reproducibility was ensured by using a fixed random seed (seed = 1). All analyses and visualizations were performed in R using the packages R, readr, dplyr, igraph, and ggplot2 (9, 20-23).

## Results

### Structural Prediction and Domain Extraction of ALPs

We annotated the structures of newly identified ALPs by homology modeling, using the classification scheme of previously curated *Mbb. smithii* and *Msp. stadtmanae* ALP dataset as a reference (14). Consistent with our earlier observations, the newly described ALPs also comprise linear strings of repeating domains. Visual inspection revealed the same four principal domain categories: the MAD (Membrane Anchor Domain), ABD (Archaeal Big Domain), RBH (Repeat-Binding Helix) domain, and a group of miscellaneous accessory domains, such as transglutaminase or pseudomurein-binding protein, that did not fit the other domain types, collectively referred to as “Others.” Attempts to map these architectures using traditional sequence-based methodologies, such as BLAST, proved insufficient to assign modular domain architectures. The high degree of sequence variability across large protein segments resulted in poor sensitivity and specificity, failing to capture the underlying structural organization.

To address these technical hurdles, we used PRISM (Protein Recognition and Integrated Segmentation Model), which utilizes a deep-learning architecture that interprets the underlying structural topology and spatial patterns of these proteins, PRISM bypasses the constraints of sequence-based comparisons (24). Using PRISM and manual curation, we generated a set of 773 ALPs from 17 different *Methanobacteriales* species. The results show that certain architecture types, e.g. Group 1 and Group 2, appear to be most common for ALPs across most analysed species, while ALPs with 1 or more “other” domains also constitute a significant proportion of the ALP repertoire (Fig1A and Table S2). The overall frequency of RBH and “other” domains per ALP was low when compared to that of ABDs, which were present with more than 40 repeats in some ALPs. A sequence-based comparative analysis revealed that some ALPs appear to share similarity with other ALPs within species, however, similarity of some ALPs across species or genera could also be observed. Likewise, a clustering by environment could be observed for some smaller clades, particularly for thermophilic environmental species (Fig 2).

**Figure 1.**
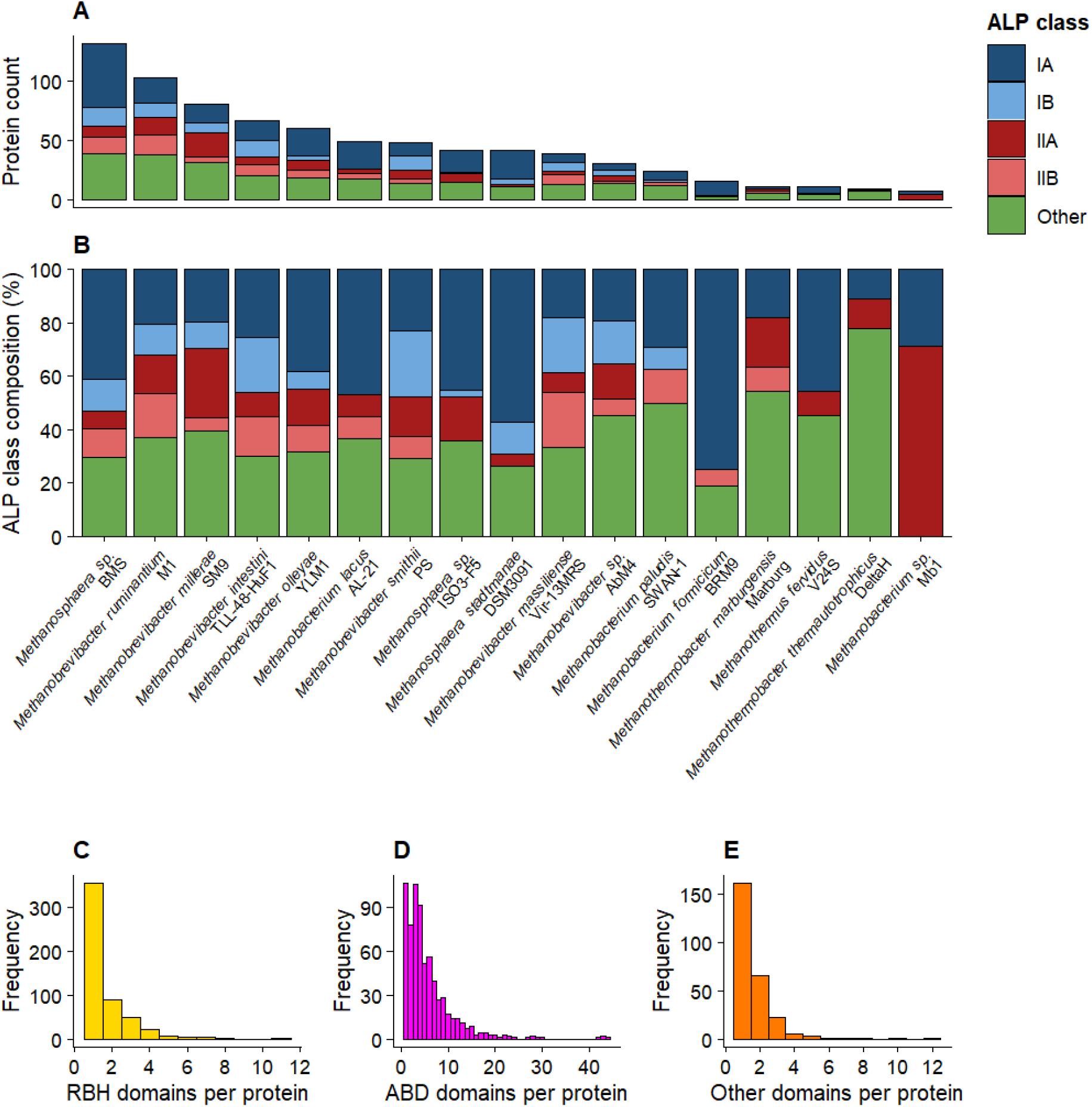
Common domain architectures in ALPs from *Methanobacteriales* strains. Shown are the absolute number of ALP groups per organism (A), the relative number of groups per organisms (B), and the frequency of ABD, RBH, and ‘Other’ domains detected in all ALPs from the 17 species that were analysed.

**Figure 2.**
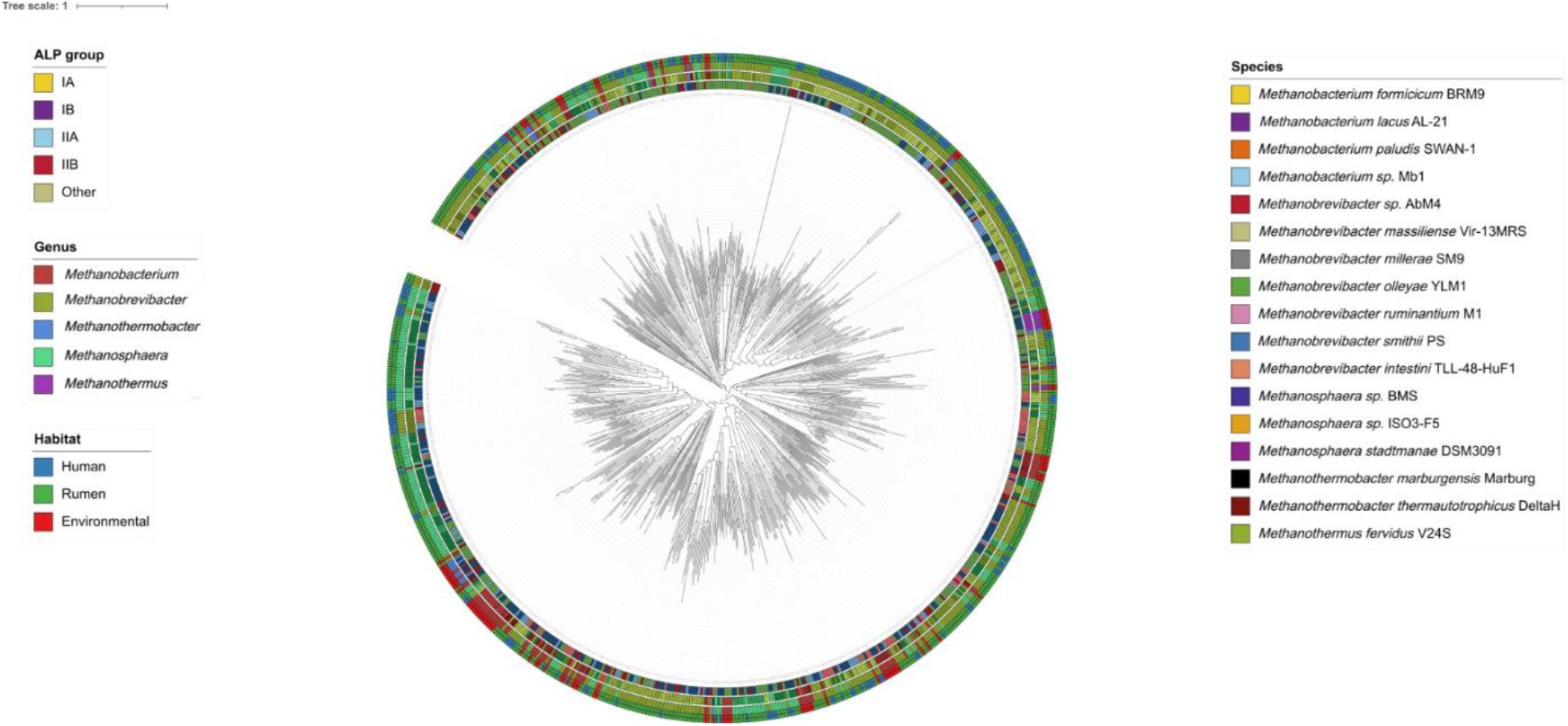
Phylogenetic distribution and structural classification of Adhesin-Like Proteins (ALPs) *Methanobacteriales* species. This circular maximum-likelihood tree illustrates the evolutionary divergence of ALPs, with branch lengths representing evolutionary distance measured in expected substitutions per site, as determined by FastTree. The metadata rings surrounding the tree provide structural, taxonomic, and ecological context for each sequence and are numbered from the centre outward. The first (innermost) ring categorizes proteins into ALP Groups based on the specific arrangement of Archaeal Big Domains (ABDs) and right-handed beta-helical domains (RBHs). Group IA consists of sequences in which all RBHs precede all ABDs in the protein sequence, whereas Group IB contains proteins with interspersed arrangements of these domains. Group II comprises proteins containing only ABDs (IIA) or only RBHs (IIB), while the “Other” category includes proteins containing domains in addition to the standard ABD and RBH domains. The second and third rings denote the species and genus of the source organism, respectively, and the fourth (outermost) ring indicates the ecological niche of the species, distinguishing between the human gut, rumen, and various environmental habitats.

### ALPs constitute a substantial and variable fraction of *Methanobacteriales* genomes

Previous analyses of individual methanogen genomes indicated the presence of ALPs in most *Methanobacteriales* genomes, however, a comparative analysis of ALPs across multiple genera of this order had not been performed previously. Our analysis show that the number of ALPs may vary substantially between representative member of the *Methanobacteriales* (Fig 3). Across the analysed methanogen genomes, ALPs represented a considerable proportion of the gene repertoire, ranging from low (*Methanobacterium* sp. Mb1= 0.4 %) to relatively high percentage (*Methanosphaera* sp. BMS= 6.3%) depending on the species (Fig. 3A). While the overall trend was consistent when expressed relative to genome size, notable interspecies variation was observed, indicating differential expansion of ALP-encoding genes across lineages. The relative abundance of ALPs calculated as a percentage of genes closely mirrored values normalized to genome size (Fig. 3A), suggesting that differences in ALP content are not solely driven by genome size variation but reflect genuine differences in gene allocation. ALP lengths varied widely both within and between species (Fig.3B). Some species exhibited relatively narrow distributions, whereas others showed broad ranges spanning several hundred to thousands of amino acids, consistent with complex domain architectures. The number of ALPs per species differed substantially (Fig. 3B), with some genomes encoding large repertoires of ALPs (e.g. *Methanosphaera* BMS = 135), while other species, such as *Methanobacterium* sp. MB1, harboured fewer than 10 ALPs. This expansion was not strictly correlated with protein length distributions, suggesting that both the number and structural diversity of ALPs contribute to species-specific functional potential. Noteworthy in this regard is also the genome of *Methanosphaera* sp. ISO3-F5, which encodes 46 ALPs, while the mean ALP length was the highest among all analysed strains (∼2020 amino acids).

**Figure 3.**
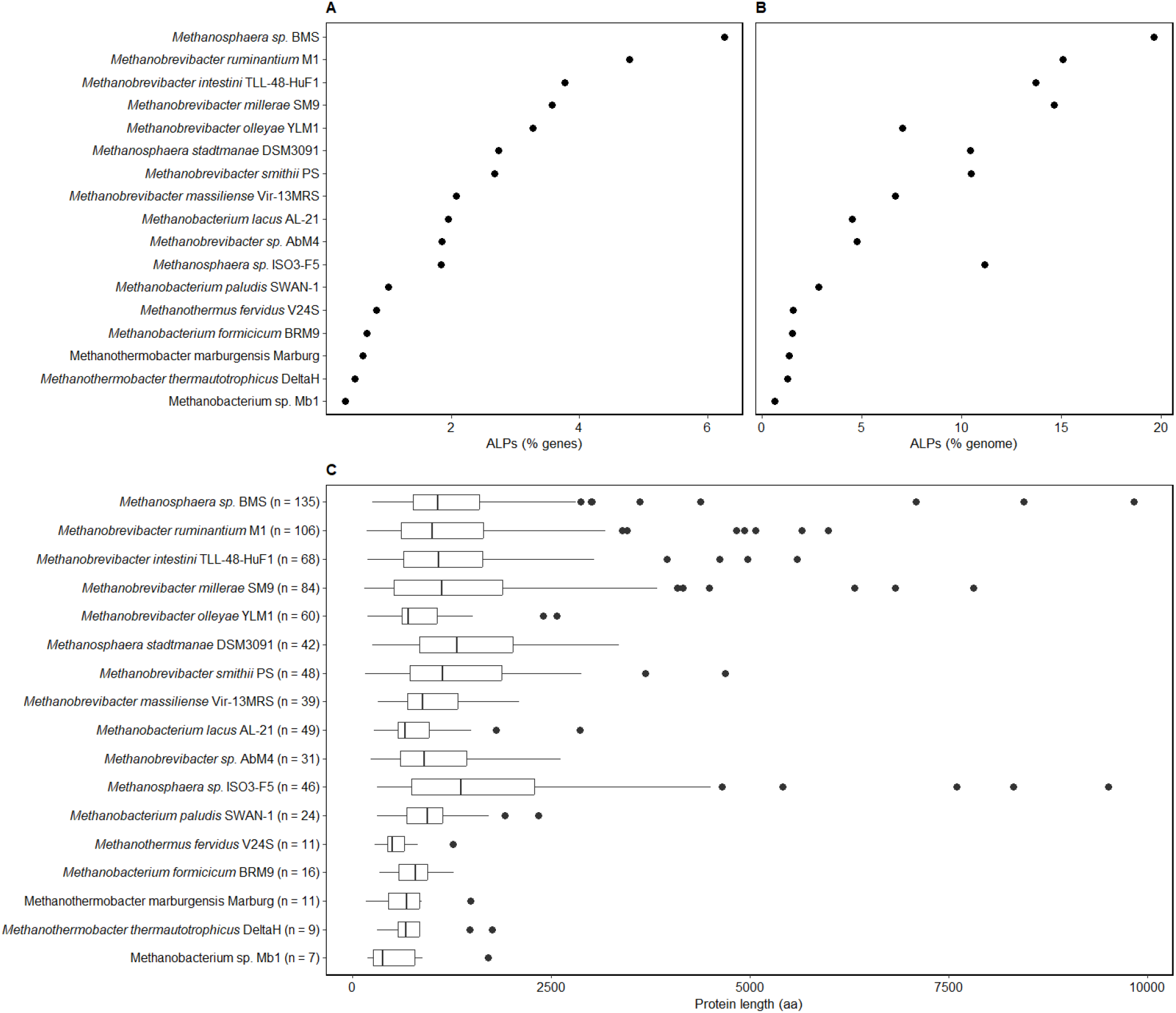
(A) The proportion of ALP-encoding genes relative to the total number of predicted protein-coding genes (% genes) and (B) genome size (% genome) across methanogen species. Species are ordered by the percentage of ALPs relative to total gene count. (C) Distribution of ALP protein lengths for each species. Boxplots show the median and interquartile range, with whiskers indicating 1.5× the interquartile range; points represent outliers. The number of ALPs identified per species is indicated in parentheses.

### Species-specific expansion of ALP repertoires in *Mbb. smithii* and *Mbb. intestini*

Our results indicate that ALP number and length varied substantially between species, however, as only one representative genome per strain was analysed it could also be argued that the variation is not species-specific but rather strain specific. As high-quality /closed genomes are currently not available for most strains of the same *Methanobacteriales* species, we extended this analysis to just two closely related species, *Mbb. smithii* and *Mbb. intestini*, where the later has only recently been described as a species (25). It was observed that the total number of ALPs did vary significantly between the two species (Wilcoxon rank sum, p=0.00295), while the length of the ALPs between the two species did not appear to differ significantly (p=0.606). While variation could be observed, both of number and length, this finding may indicate that these features are characteristic of a species-specific ALP repertoire (Fig 4).

**Figure 4.**
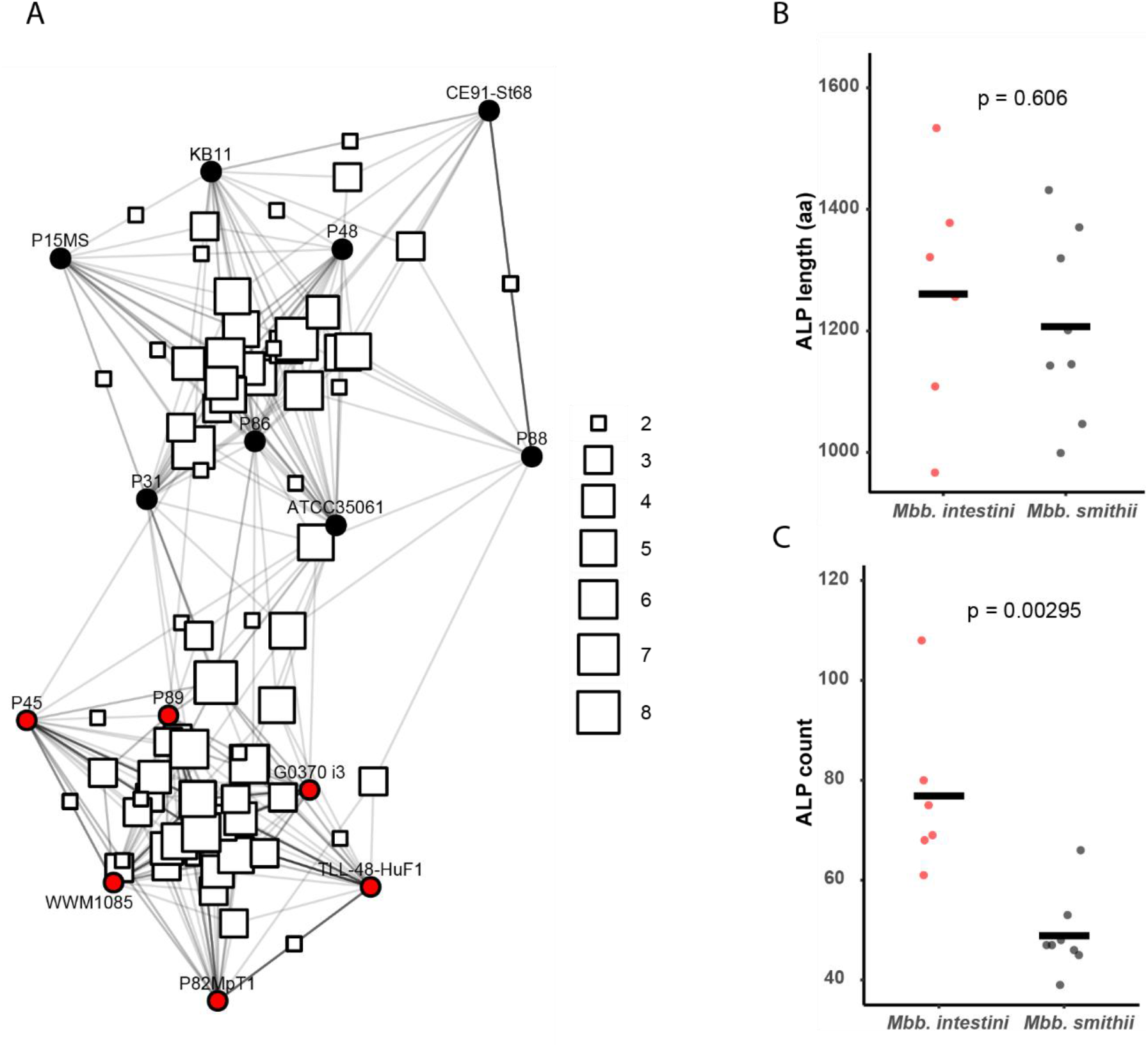
Comparative sequence-based analysis of ALPs from *M. smithii* and *M. intestini* strains. (A) Network analysis of ALPs shared between *M. smithii* (black) and *M. intestini* (red) strains. (B) mean length variation of ALPs from the two species. (C) ALP count of strains for each of the species.

Applying stringent similarity cut-offs for ALP length and sequence identity, we also performed a sequence-based comparative analysis to determine which ALPs may be conserved across both species and/or just within a species. A network analysis of ALPs from these two species revealed that ALPs appear to be more commonly shared among strains of the same species (Fig 4). Our results indicate that even within a species, relatively few ALPs are shared among all strains and while sharing of similar ALPs across species is observed, it did not seem to be the case that an ALP was shared among all strains of both species. It is noteworthy that sequence dissimilarity between species may make it difficult to identify ALPs that are shared among different species. We did note that species may harbor similar ALPs, when considering the domain architecture of the ALPs, despite showing limited sequence similarity (Fig 5). Examples are clusters S15A and S15B, which appear to be species-specific clusters that show limited sequence similarity, but that have the same architecture (3 ABDs). Other example would be S33A and S34A, which has a domain architecture that is conserved in 12 out of 14 strains. Overall, smaller ALPs with less complex architectures were more frequently detected across multiple species and environments (Fig. S1 and Fig. S2), while a substantial proportion of ALPs appeared to be strain-specific.

**Figure 5.**
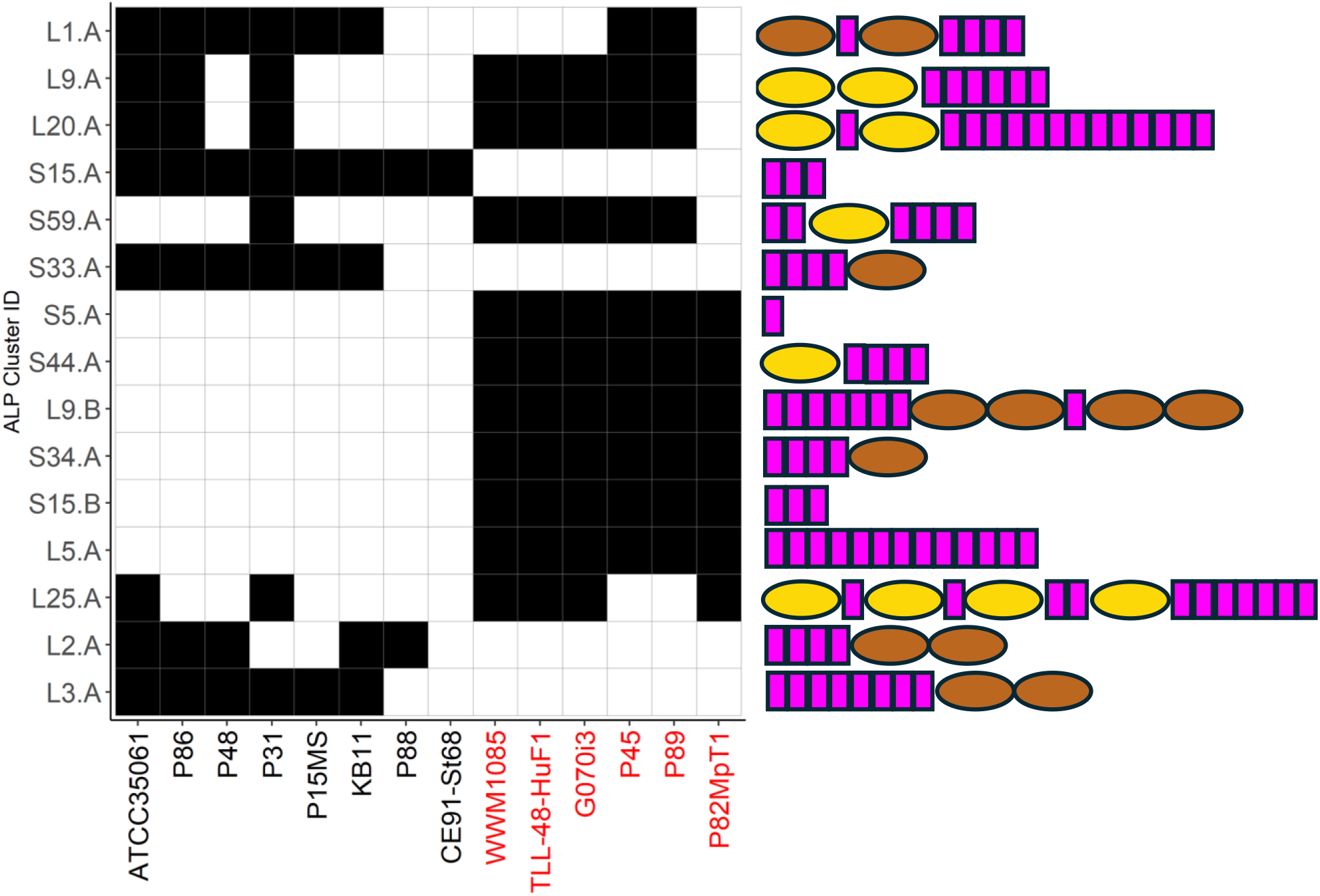
Domain composition of ALPs shared between *Mbb. intestini* and *Mbb. smithii*. Only ALPs that are found in at least six strains of the two species combined are depicted. Strain IDs are shown on the x-axis; *Mbb. smithii* strain names are shown in black and *Mbb. intestini* strain names in red. ABDs are shown in magenta, RBHs in yellow, and other domains in brown. Domains are schematic and sizes are not drawn to scale.

## Discussion

In this study, we showed that the same ALP-associated domains can be observed across a broad range of species within the order *Methanobacteriales*. While domain architectures vary considerably within individual strains, thereby giving rise to a diverse repertoire of ALPs, we also observed that the recently introduced schematic ALP grouping established for *Mbb. smithii* and *Msp. stadtmanae* can likewise be applied to additional members of this order. It should, however, be emphasized that this grouping does not necessarily imply functional conservation, but instead primarily serves to categorize ALPs according to their domain architecture. Even within individual groups, such as group IA, substantial variation can be observed, particularly with respect to the copy number of certain domain types. This is especially apparent for ABDs, where copy numbers can range widely (n = 1–44). The functional significance of these varying copy numbers remains speculative at present, but it is conceivable that copy numbers, and consequently overall ALP lengths, may be associated with distinct ALP functions, similar to what may be hypothesized for the broader domain architectures.

Likewise, ALPs containing “Other” domains appear to represent one of the more prevalent ALP groups. The annotation of many of these domains remains vague and will require experimental validation. Nevertheless, the presence of one of the more frequently observed domains, the pseudomurein-binding domain (PBD), may point toward interactions involving methanogens that possess pseudomurein-containing cell walls, as is characteristic of members of the *Methanobacteriales*. This could potentially indicate cell–cell interactions either between different methanogen species or within a single species, thereby contributing to adhesion processes and/or types of biofilm formation. In this context, it is also noteworthy that a recent study reported that small extracellular vesicles secreted by *Mbb. smithii* and *Msp. stadtmanae* in monoculture appeared to be enriched in ALPs (22, 26). A more detailed analysis of these ALPs may therefore provide important insights into both the general biological role of ALPs and the specific functions associated with individual ALP domains.

Our results reveal that *Mbb*. species, particularly *Mbb. smithii* and *Mbb. intestini*, may harbor ALP repertoires with species-specific characteristics. For example, a statistically significant difference in ALP numbers was observed between the two species. This difference could point toward species-specific adaptations within the gut environment and warrants further investigation, particularly given that these taxa have only relatively recently been recognized as distinct species (25). While ALP repertoires generally appear to be species-specific, the presence or absence of conserved ALPs across different species may also reflect ecological adaptations of *Methanobrevibacter* species to distinct environmental niches.

Expanding these analyses to include a larger number of high-quality genomes, together with the continued improvement of structure prediction and functional annotation approaches, would enable broader comparative analyses across additional species and strains. Genomes reconstructed from metagenomic sequencing datasets could likewise be incorporated, provided that the assemblies are sufficiently complete, consist of only a limited number of contigs, and meet high-quality standards. However, this remains challenging because metagenomic sequencing typically favours the recovery of the more abundant genomes within a sample, whereas methanogens often constitute only a minor proportion of the microbiome. The application of deep sequencing and hybrid-assembly strategies may help to overcome some of these limitations, as has recently been demonstrated in human cohort studies (23). Nevertheless, even in those studies, contiguous methanogen assemblies were primarily obtained from samples with relatively high methanogen abundance (27).

This study focuses on the order *Methanobacteriales*. However, it should be noted that ALPs have also been reported in other methanogenic orders, particularly the *Methanomassiliicoccales* (20, 21). Although the number of fully sequenced genomes available for these methanogens remains comparatively limited, preliminary analyses suggest that their ALPs may share certain common features with those of the *Methanobacteriales*, while also exhibiting distinct characteristics that warrant separate and more in-depth investigation. Given that members of the *Methanomassiliicoccales* are also important constituents of the rumen and other intestinal environments, defining the ALP repertoire of this order will be essential for a broader understanding of ALP diversity and function in gut-associated methanogens.

Although this study focuses on methanogens originating from the mammalian intestinal tract, members of the *Methanobacteriales* (and *Methanomassiliicoccales*) are widely distributed across diverse host-associated and environmental ecosystems, including the gastrointestinal tracts of insects (28).. Draft genomes from insect-associated *Methanobrevibacter* species and wet-wood-associated methanogens suggest the presence of ALP-encoding genes in these organisms as well (29, 30). However, although several high-quality genomes from insect-associated methanogens have recently become available, broader taxonomic and ecological sampling will be required for a comprehensive comparison of ALPs across the full diversity of *Methanobacteriales* (28). In conclusion, it can be stated that the novel approach for ALP domain detection does allow a rapid and accurate annotation of ALPs from methanogen genomes and sheds light on the diversity of domains and domain-architecture across different species. Rate-limiting steps for the annotation are currently the availability of high-quality genomes and also the structural prediction of ALPs. Both can be overcome with the use of improved sequencing approaches, e.g. including long-read sequencing and deep sequencing of genomes/enrichment cultures, and with increased use of computational resources. Our results highlight that adhesiomes are abundant across many different taxa and appear to possess species-specific characteristics. Further research into the function and molecular mechanisms of ALPs will be necessary to better understand their role in speciation and the occupation of ecological niches.

## Author contributions

AT, ABG, SR and HS conceived the study. AT, ABG, SR and HS analyzed the data. AT and HS wrote revised the manuscript. ABG and SE reviewed and edited the manuscript. HS organized and supervised the study.

## Funding

This research was supported by core funding from Temasek Lifesciences Laboratory

## Conflicts of interest

The authors declare that they have no competing interests.

## Acknowledgments

We thank the TLL IT department for their support with computational resources. We also acknowledge the use of ChatGPT (https://www.chatgpt.com/) to improve the grammar and wording of the manuscript.

## Notes

### Competing Interest Statement

The authors have declared no competing interest.

