## Supplementary material for "Comparative analysis of adhesin-like proteins from *Methanobacteriales* species": Table S1

Table S1. Metadata on strains and ALPs

| Genus | Species | Strain | Source/ accession number |
|---|---|---|---|
| *Methanothermobacter* | *marburgensis* | Marburg | CP001710.1 |
| *Methanosphaera* | *stadtmanae* | DSM3091 | CP000102.1 |
| *Methanobrevibacter* | *sp.* AbM4 | | CP004050.1 |
| *Methanosphaera* | *sp.* BMS | | CP014213.1 |
| *Methanothermus* | *fervidus* | DSM2088 | CP002278.1 |
| *Methanobacterium* | *formicicum* | BRM9 | CP006933.1 |
| *Methanosphaera* | *sp.* ISO3-F5 | | CP118753.2 |
| *Methanobacterium* | *lacus* | AL-21 | CP002551.1 |
| *Candidatus* Methanobrevibacter | *massiliense* | Vir-13MRS | JARBXM010000000 |
| *Methanobacterium* | *sp.* MB1 | | HG425166 |
| *Methanobrevibacter* | *millerae* | SM9 | CP011266.1 |
| *Methanobrevibacter* | *olleyae* | YLM1 | CP014265.1 |
| *Methanobacterium* | *paludis* | SWAN1 | CP002772.1 |
| *Methanobrevibacter* | *ruminantium* | M1 | CP001719.1 |
| *Methanothermobacter* | *thermautotrophicus* | Delta H | AE000666.1 |
| *Methanobrevibacter* | *smithii* | P15MS | https://github.com/SDMUG/Methanobrevibacter_Enrichment/blob/main/Nanopore%20Sequencing/contigs/P15MS_consensus.fasta |
| *Methanobrevibacter* | *smithii* | P31 | https://github.com/SDMUG/Methanobrevibacter_Enrichment/blob/main/Nanopore%20Sequencing/contigs/P31_consensus.fasta |
| *Methanobrevibacter* | *smithii* | P48 | https://github.com/SDMUG/Methanobrevibacter_Enrichment/blob/main/Nanopore%20Sequencing/contigs/P48_consensus.fasta |
| *Methanobrevibacter* | *smithii* | P86 BT | https://github.com/SDMUG/Methanobrevibacter_Enrichment/blob/main/Nanopore%20Sequencing/contigs/P86_BT_consensus.fasta |
| *Methanobrevibacter* | *smithii* | P88 | https://github.com/SDMUG/Methanobrevibacter_Enrichment/blob/main/Nanopore%20Sequencing/contigs/P88_consensus.fasta |
| *Methanobrevibacter* | *smithii* | KB11 | CP017803. |
| *Methanobrevibacter* | *smithii* | CE91-St68 | AP025587.1 |
| *Methanobrevibacter* | *smithii* | ATCC 35061 | CP000678 |
| *Methanobrevibacter* | *intestini* | WWM1085 | NQLD00000000 |
| *Methanobrevibacter* | *intestini* | TLL-48-HuF1 | CP081485. |
| *Methanobrevibacter* | *intestini* | P89 | https://github.com/SDMUG/Methanobrevibacter_Enrichment/blob/main/Nanopore%20Sequencing/contigs/P89_consensus.fasta |
| *Methanobrevibacter* | *intestini* | P82MpT1 | https://github.com/SDMUG/Methanobrevibacter_Enrichment/blob/main/Nanopore%20Sequencing/contigs/P82MpT1_sans_C_consensus.fasta |
| *Methanobrevibacter* | *intestini* | P45 | https://github.com/SDMUG/Methanobrevibacter_Enrichment/blob/main/Nanopore%20Sequencing/contigs/P45_consensus.fasta |
| *Methanobrevibacter* | *intestini* | G0370 i3 | CP187956.1 |
