## Supplementary material for "Comparative analysis of adhesin-like proteins from *Methanobacteriales* species": Table S2

Table S2. ALP and ALP class distribution in genomes

| Species | genome size | total protein-coding genes | percentage of genes | percentage of genome | ALP class IA | ALP class IB | ALP class IIA | ALP class IIB | ALP class Other |
|---|---|---|---|---|---|---|---|---|---|
| *Methanobacterium lacus* AL-21 | 2583753 | 2512 | 2.0 | 4.5 | 46.9 | 0.0 | 8.2 | 8.2 | 18 |
| *Methanobacterium formicicum* BRM9 | 2449988 | 2352 | 0.7 | 1.5 | 75.0 | 0.0 | 0.0 | 6.3 | 3 |
| *Methanobacterium paludis* SWAN-1 | 2546541 | 2339 | 1.0 | 2.8 | 29.2 | 8.3 | 0.0 | 12.5 | 12 |
| *Methanobacterium sp* . Mb1 | 2029766 | 2021 | 0.3 | 0.6 | 28.6 | 0.0 | 71.4 | 0.0 | 0 |
| *Methanobrevibacter intestini* TLL-48-HuF1 | 1951395 | 1777 | 3.8 | 13.7 | 25.4 | 20.9 | 9.0 | 14.9 | 20 |
| *Methanobrevibacter massiliense* Vir-13MRS | 1834388 | 1870 | 2.1 | 6.7 | 17.9 | 20.5 | 7.7 | 20.5 | 13 |
| *Methanobrevibacter millerae* SM9 | 2543538 | 2269 | 3.6 | 14.6 | 19.8 | 9.9 | 25.9 | 4.9 | 32 |
| *Methanobrevibacter olleyae* YLM1 | 2201192 | 1834 | 3.3 | 7.0 | 38.3 | 6.7 | 13.3 | 10.0 | 19 |
| *Methanobrevibacter ruminantium* M1 | 2937203 | 2154 | 4.8 | 15.1 | 20.4 | 11.7 | 14.6 | 16.5 | 38 |
| *Methanobrevibacter smithii* PS | 1853160 | 1795 | 2.7 | 10.5 | 22.9 | 25.0 | 14.6 | 8.3 | 14 |
| *Methanobrevibacter sp* . AbM4 | 1998189 | 1671 | 1.9 | 4.7 | 19.4 | 16.1 | 12.9 | 6.5 | 14 |
| *Methanosphaera sp* . BMS | 2868093 | 2108 | 6.3 | 19.6 | 40.9 | 12.1 | 6.8 | 10.6 | 39 |
| *Methanosphaera sp* . ISO3-F5 | 2506044 | 2281 | 1.8 | 11.2 | 45.2 | 2.4 | 16.7 | 0.0 | 15 |
| *Methanosphaera stadtmanae* DSM3091 | 1767403 | 1534 | 2.7 | 10.4 | 57.1 | 11.9 | 4.8 | 0.0 | 11 |
| *Methanothermobacter marburgensis* Marburg | 1639135 | 1752 | 0.6 | 1.4 | 18.2 | 0.0 | 18.2 | 9.1 | 6 |
| *Methanothermobacter thermautotrophicus* DeltaH | 1751377 | 1802 | 0.5 | 1.3 | 11.1 | 0.0 | 11.1 | 0.0 | 7 |
| *Methanothermus fervidus* V24S | 1243342 | 1311 | 0.8 | 1.6 | 45.5 | 0.0 | 9.1 | 0.0 | 5 |
