## supporting figures for "Comparative analysis of adhesin-like proteins from *Methanobacteriales* species"

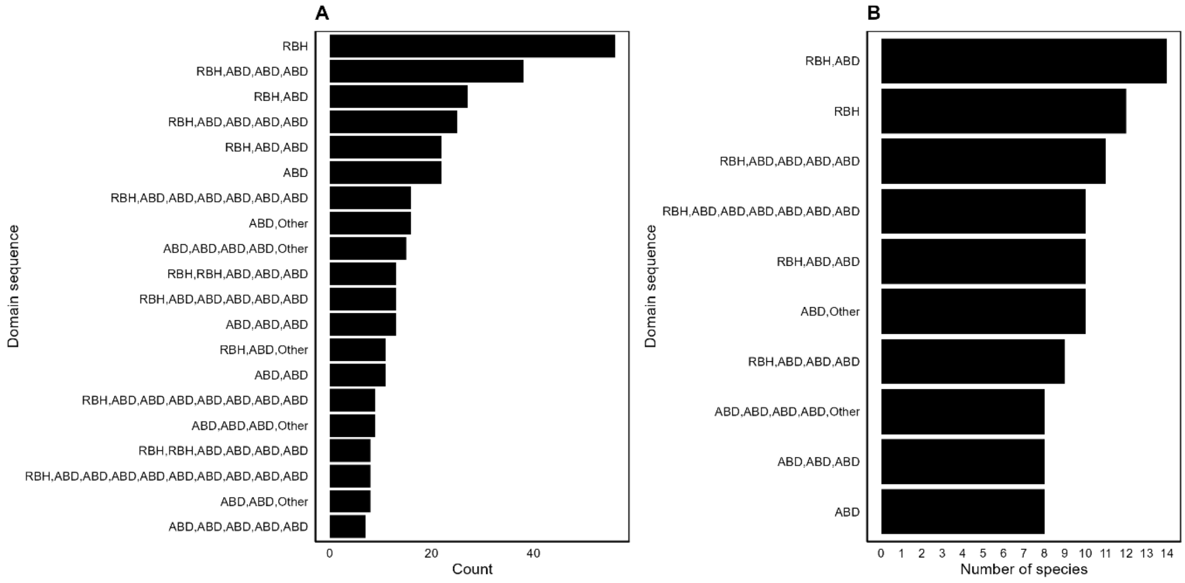


**Figure S1. Distribution and conservation of domain architectures in adhesin-like proteins (ALPs).** (A) The 20 most abundant domain sequences across all analysed proteins. In panel B, each domain sequence was counted once per species to assess its distribution across taxa. Together, these analyses highlight both the most frequent and the most widely conserved ALP architectures. Only the 10 domain sequences present in the highest number of species are shown.

### Figure S2.

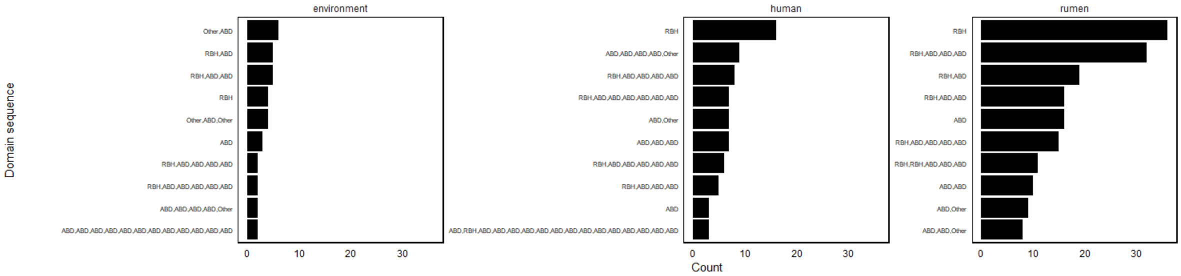


**Figure S2.** Environment-associated distribution patterns of adhesin-like protein (ALP) domain architectures. The 10 most abundant domain sequences identified within each environment.
